# Dopaminergic neuronal dysfunction induced by newer generation systemic insecticides in *Caenorhabditis elegans*

**DOI:** 10.64898/2026.06.14.732178

**Authors:** Adam R Filipowicz, Keane Bui, Nada Osman, Katherine S Morton, Isabel W Kenny-Ganzert, David R Sherwood, Joel N Meyer, Patrick Allard

## Abstract

While a growing number of studies have linked environmental exposures and Parkinson’s disease (PD)^1–3^, the impact of many pesticides remains understudied^4,5^; for example, neonicotinoids are the most used insecticides in the world, but research into their contribution to PD is limited to a handful of studies^6–8^. Newer pesticides, such as the butenolide flupyradifurone (FPF), specifically developed to overcome increased pest resistance^9^ and spurred on by tighter restrictions on neonicotinoids such as imidacloprid (IMI)^10^, are even less studied. New approach methodologies (NAMs) that allow for rapid evaluation of pesticide exposures are needed to evaluate potential links between the growing number of pesticides and PD^11^. To this end, we exposed the model nematode *Caenorhabditis elegans*^12^ to IMI and FPF. Due to its high degree of tractability, and conservation of many genetic, neuronal, and toxic mode of action processes, *C. elegans* has been invaluable in both elucidating mechanisms and novel therapeutic targets for PD that can be validated in other models^13^, and as a complementary tool for early toxicity screening^14^. Along this line, we found that exposure to IMI, and to a greater extent FPF, in young adult animals causes significant dendritic blebbing, an early sign of neurodegeneration, exclusively in dopaminergic neurons. Blebbing was accompanied by impairment of dopamine-mediated behaviors, changes in neuronal mitochondrial morphology, and elevation of pathways related to reactive oxygen species (ROS). We were able to reduce the blebbing caused by IMI and FPF two ways: 1) pharmacologically via administration of the antioxidant N-acetyl cysteine (NAC); 2) genetically via knockout of a MAP kinase (MAPK) stress response pathway. This suggests that oxidative stress is a key mediator of this insecticide-induced dopaminergic neurodegeneration.

## Results and Discussion

Parkinson’s disease (PD) is the most common neurodegenerative movement disorder and second-most common neurodegenerative disorder, affecting around 1-3% of the population over 65 years of age^15^. Degeneration of dopaminergic neurons in the substantia nigra, along with intracellular α-synuclein accumulation, are hallmarks of the disease^16^. While numerous studies have sought to determine genetic^17–20^ and environmental PD risk factors^1–3^, studies on the latter are largely associative. The estimated heritability of PD is only 0.34, meaning the significant majority of the risk for developing idiopathic PD is attributable to environmental factors^21^. Mechanistic work on environmental risk factors has mainly focused on a small number of pesticides such as paraquat and rotenone^22–26^. This leaves many pesticides, including some of the most widely used, understudied. Neonicotinoids, for example, account for approximately 30% of the global insecticides market^27^, and although several studies have shown that neonicotinoids induce adverse neurobehavioral effects in nontarget organisms^28,29^, little research has been done on their potential role in human neurodegenerative diseases such as PD. Neonicotinoids are nicotinic acetylcholine receptor (nAChR) agonists with higher binding affinities for insect nAChRs than mammalian receptors^30^. Nevertheless, this lower affinity might still be problematic, as recent studies using a human dopaminergic cell line have shown that imidacloprid (IMI) and other neonicotinoids are able to activate intracellular calcium signaling through nAChRs at low micromolar concentrations^7^. Sustained cholinergic activation could lead to excitotoxicity and oxidative stress^31^. Furthermore, IMI has significant effects on protein synthesis and ribosomal function in this cell line^8^.

As pests become resistant to neonicotinoids and their use lessens due to stricter regulations other nAChR agonists, such as the novel butenolide flupyradifurone (FPF), have come onto the market^9,10^. Little is known about the potential off-target neurotoxic effects of FPF, although an increasing number of reports indicate adverse effects, including lethality and abnormal behaviors in bees and other non-pest animals^32–34^, and its long environmental half-life suggests significant potential for exposure^35^. To investigate nAChR agonist-mediated dopaminergic neurotoxicity and uncover potential mechanisms related to PD, we exposed the model nematode *C. elegans* to IMI and FPF. *C. elegans* is a highly relevant and widely used model of PD^13^. The nematode’s 8 dopaminergic neurons are easily visualized using fluorescent reporter strains that are used to investigate neurodegenerative phenotypes including axonal and dendritic blebbing and to screen for neurotoxicants and neuroprotective effects in the context of PD^36–38^. Well studied mammalian dopaminergic neurotoxicants, including MPP+ and 6-OHDA, induce dopaminergic neurodegeneration in *C. elegans,* and work in the nematode has uncovered the neurodegenerative potential of manganese, methylmercury, rotenone, and paraquat^13,36,38–42^. In addition to these morphological phenotypes, behaviors that are directly dependent on dopaminergic circuits have also been extensively used to identify disease modifiers. These include basal slowing^43^, or the reduction in movement in the presence of food, and foraging^44^. *C. elegans* dopaminergic neurons are also sensitive to overexpression of well-characterized human pro-PD factors such as α-synuclein and LRRK2, which cause a loss of dopamine neurons and deficits in dopamine dependent behavior^13,45,46^. In addition, genetic loss-of-function models of PD have been highly studied in *C. elegans,* including mutations in the *C. elegans* homologs of PRKN, PINK1, DJ-1, and ATP13A2, and have been combined with environmental exposures to further our understanding of gene-environment interactions in PD^47,48^. We therefore applied this powerful model to first establish sublethal exposure conditions for IMI and FPF. We found that following a 48-hour exposure of L4 stage animals to IMI or FPF, all doses tested (18μM, 55μM, 166μM and 500 μM) were non-lethal (Figures 1A and 1B; N=4 exposures). Conversely, exposure to the potent nematicide methomyl^49^ caused significant lethality at 18μM (Figure S1), the lowest tested dose, demonstrating the efficacy of our exposure conditions. Six other pesticides were lethal at 500μM (chlorpyrifos, fipronil, carbofuran, mancozeb, diazinon, and malathion), while eleven others tested, including other neonicotinoids, were non-lethal at all concentrations (Figure S1). We selected IMI and FPF to follow up based largely on widespread use of IMI as the most popular neonicotinoid and FPF as a new generation nAChR agonist insecticide designed to replace neonicotinoids in case of resistance.

**Figure 1.**
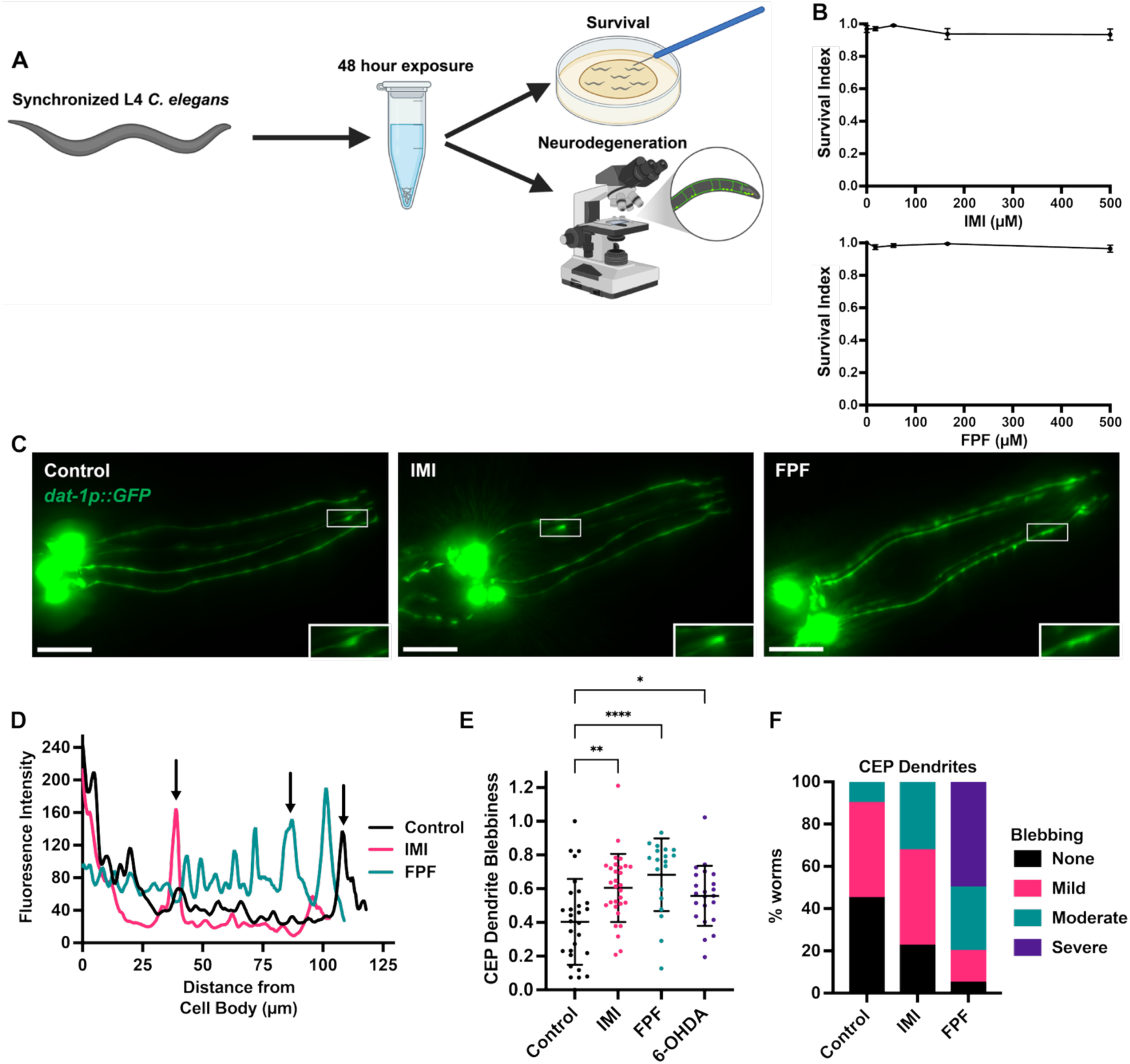
IMI and FPF induce blebbing in CEP dendrites. (A) Schematic of the pesticide exposure paradigm used in this study. Synchronized L4 animals were exposed to vehicle control (0.1% DMSO) or pesticides (18-500μM, 3x serial dilution) for 48 hours in liquid culture. Animals were then washed and either transferred to a plate to check for survival or staged on a microscopy slide for visualization of different neuronal subtypes using strains expressing GFP driven by neuron-specific promoters: *dat-1p* (dopaminergic), *unc-17p* (cholinergic), or *tph-1p* (serotonergic). (B) Survival index for IMI and FPF at 18, 55, 166, and 500μM. Dots are the mean at each concentration and error bars are SEM. N=4 exposures. (C) Representative images of *dat-1p::GFP* animals exposed to vehicle control (0.1% DMSO) or 500μM of IMI or FPF for 48 hours. Four CEP neuronal bodies and four CEP dendrites extending to the nose of the animal in the upper right are visible. Insets highlight blebs along CEP dendrites. Scale bar, 20μm. (D) Representative fluorescence intensity profile along the length of CEP dendrites from the images in (C). Arrows pointing to spikes in fluorescence correspond to bleb insets in (C). (E) Quantification of blebbiness of CEP dendrites after 48-hour exposure to vehicle control (0.1% DMSO), IMI or FPF (500μM), or 6-OHDA (10mM). See Methods section for details on blebbiness calculation. One-way ANOVA with Dunnett’s multiple comparisons test was used to determine significance. * is p<0.05, ** is p<0.01, and **** is p<0.0001. Each dot is an individual animal. The line is the mean and error bars are standard deviations. N=2-3 exposures; n=19-30 animals. (F) Blind scoring of CEP dendrites after 48-hour exposure to vehicle control (0.1% DMSO) or IMI/FPF (500μM). Two blinded scorers determined the level of blebbing using the following definitions: None: no blebs observed, Mild: at least one bleb on one dendrite, Moderate: multiple blebs on at least one dendrite, Severe: multiple blebs on multiple dendrites. N=2-3 exposures; n=19-30 animals.

Previous reports have shown that dopaminergic neurotoxicants cause breakages and blebbing (protrusions from the string-like dendrites) that are especially evident on the cephalic (CEP) dendrites^50^. Using a *C. elegans* strain expressing GFP in dopaminergic neurons (*dat-1p::GFP)* we observed that after a 48-hour exposure, both IMI and FPF at 500μM caused noticeable CEP dendritic blebbing compared to vehicle control (Figure 1C). Doses as low as 18μM of either pesticide also led to significant blebbing (Figure S2). To measure levels of blebbing, CEP dendrites were traced and an intensity profile was generated for each dendrite (Figure 1D). Spikes in intensity were evident in the traces and corresponded to blebs in the dendrites (insets in Figure 1C and arrows in Figure 1D). The blebbiness of each dendrite was calculated by adapting a seasonal-trend decomposition using LOESS (STL) method to the fluorescence intensity traces. Blebbiness was averaged across animals and normalized to the control exposure. This analysis revealed significant increases in blebbiness upon exposure to either pesticide, with FPF having a stronger effect than IMI (Figure 1E). We validated our blebbiness measurement in two ways. First, we measured blebbiness after exposure to 10mM 6-OHDA, a neurotoxicant known to induce degeneration of dopamine neurons in *C. elegans*^36,40,51^. Exposure to 6-OHDA resulted in significantly elevated levels of blebbing compared to control, though slightly lower than either pesticide (Figure 1E). Second, blind scoring for blebbing was performed, with animals assigned to one of four blebbing categories: none, mild (at least one bleb), moderate (a few blebs on multiple dendrites), and severe (multiple blebs on all dendrites). Some mild blebbing was noted in control animals, usually observable as a bleb near the terminus of a CEP dendrite (Figure 1C). IMI increased the number of animals displaying moderate blebbing, and half of the FPF exposed animals displayed severe blebbing (Figure 1F). Contrary to a previous study on IMI^52^, which used different exposure conditions and a commercial formulation containing IMI, IMI and FPF had no effect on serotonergic or cholinergic neurons (Figure S3). This suggests that dopaminergic neurons are particularly sensitive to these pesticides in *C. elegans*. Exposure to the pesticides during development (L1 to L4 larval stages; 48 hours) had no effect on CEP dendritic blebbing (Figure S4), indicating that the L4 to young adult transition period is particularly sensitive to IMI and FPF-induced dopaminergic neurodegeneration.

Degeneration of CEP neurons correlates with behavioral changes in exposed animals, including defects in basal slowing, odor avoidance, swimming-induced paralysis, and foraging^13,43,44,50^. These behaviors require dopamine as animals lacking *cat-2,* a tyrosine hydroxylase required for biosynthesis of dopamine, are completely defective in these activities^43,44^. We therefore tested basal slowing (slowing down in the presence of bacterial food, Figure 2A) after exposure to IMI and FPF and found that while both exposed and unexposed animals showed lower speeds on food versus off food (Figure 2B), IMI and FPF exposed animals displayed a defect in basal slowing and had significantly smaller reductions in speed (Figure 2C). To measure the impact of IMI and FPF on foraging, we conducted a food race assay (Figure 2D) and found that both pesticides also disrupted navigation to a food source (Figure 2E), with an average 50%-occupancy time for control animals of 38.02 minutes, versus 66.18 and 64.30 minutes for IMI and FPF, respectively (Figure 2F). These behavioral assays indicate that both IMI and FPF have strong and consistent deleterious effects on dopamine-mediated behaviors.

**Figure 2.**
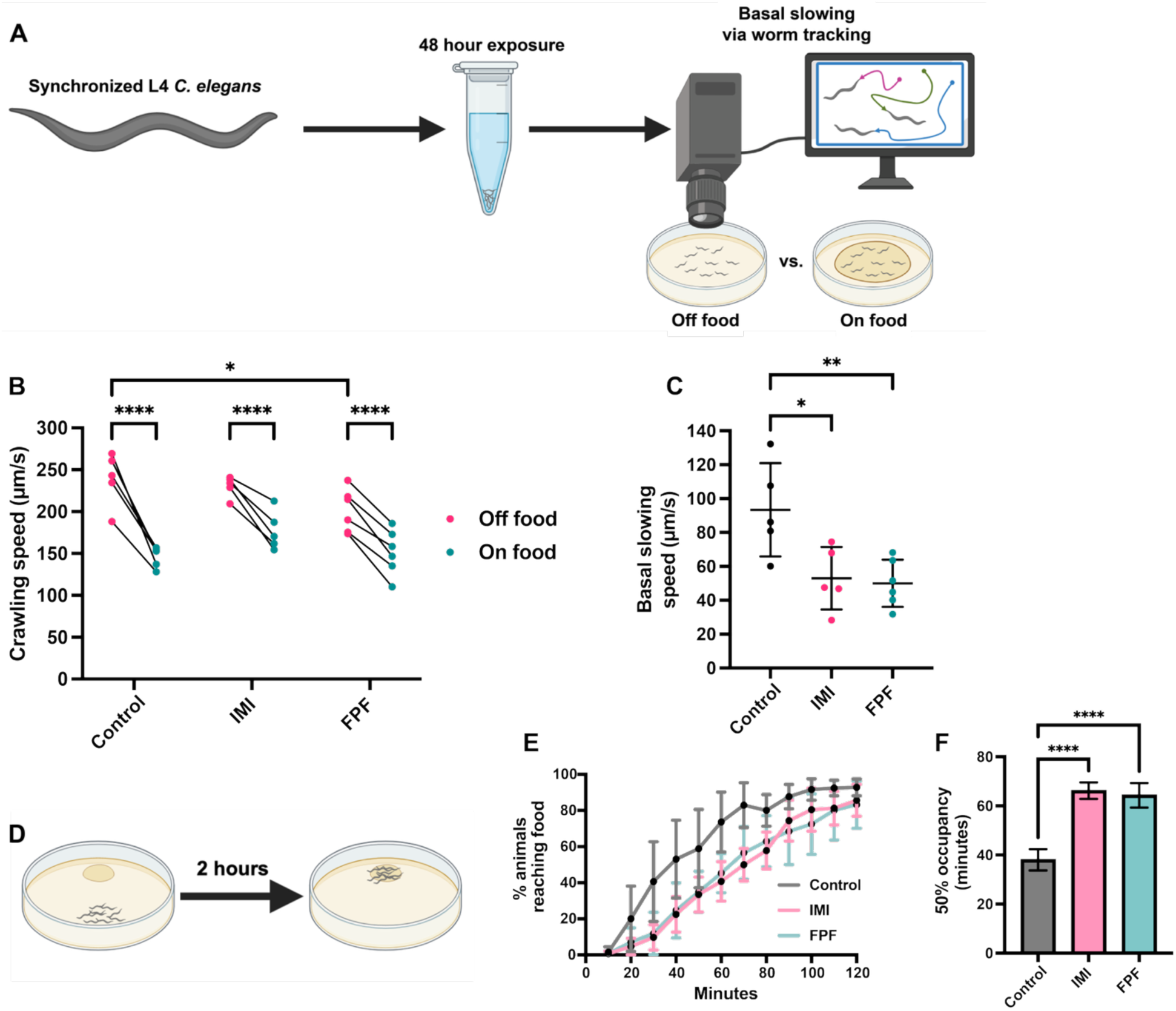
IMI and FPF disrupt dopamine-mediated behaviors. (A) Schematic of the basal slowing assay. Animals exposed to either vehicle control (0.1% DMSO) or 500μM of IMI or FPF were allowed to recover and then transferred to assay plates with or without food (a thin lawn of *E. coli* strain OP50). Animals were video recorded for 1 minute and then tracked and analyzed using WormLab. (B) Crawling speeds of paired exposures across “Off food” and “On food” groups were determined and two-way ANOVA with Dunnett’s multiple comparisons test was used to determine significance. * is p<0.05 and **** is p<0.0001. Each dot is the average speed of animals from one exposure, with lines connecting paired exposures. N=5-6 exposures. (C) Basal slowing speed was calculated as (Off food crawling speed – On food crawling speed). One-way ANOVA with Dunnett’s multiple comparisons test was used to determine significance. * is p<0.05 and ** is p<0.01. Each dot is the average speed of animals from one exposure. The line is the mean and error bars are standard deviations. N=5-6 exposures. (D) Schematic of the food race assay. Animals were exposed as in (A), recovered, and transferred to the side of a plate opposite a 20μL lawn of OP50. Animal occupancy on the lawn was recorded every 10 minutes for 2 hours. (E) Occupancy results for each exposure group expressed as percent of animals reaching the food. The mean at each time point is displayed. Error bars are standard deviation. N=4 exposures. (F) The time it took half the animals to occupy the lawn was determined using a variable slope least squares fit of the curves shown in (E). The mean of each group is displayed. Error bars are standard deviation. One-way ANOVA with Dunnett’s multiple comparisons test was used to determine significance. **** is p<0.0001. N=4 exposures.

To uncover the mechanism behind IMI and FPF induced dopaminergic neurodegeneration, we examined changes in two common and interconnected drivers of neurodegeneration: mitochondrial dysfunction and reactive oxygen species (ROS). Mitochondrial dysfunction has been implicated in both PD pathogenesis and animal models of pesticide exposure, as evidenced by MPP+ and rotenone’s ability to target complex I of the mitochondrial electron transport chain and α-synuclein accumulation in mitochondria^23,39,40,53–55^. Mitochondria respond to stress via multiple surveillance mechanisms, including the mitochondrial unfolded protein response (UPRmt) and changes in mitochondrial fission and fusion dynamics^56^. Mitochondrial fusion allows damaged mitochondrion to fuse with healthy mitochondria to restore function. However, if stress is too high then mitochondria stop fusion to limit damage. In contrast fission allows damaged mitochondria to be segregated and turned over through mitophagy^56^. To measure activation of UPRmt after IMI or FPF exposure, we exposed *hsp-6p::GFP* animals and measured whole body fluorescence. HSP-6 is a mitochondrial chaperone protein and is highly expressed upon UPRmt induction^57^. Imaging revealed a significant increase in *hsp-6p::GFP* fluorescence after 48-hour exposure to FPF, but not IMI (Figures 3A and 3B). To investigate whether IMI or FPF lead to changes in mitochondrial fission and fusion dynamics, we used a *C. elegans* strain expressing the fluorescent protein mKate2 exclusively in the mitochondria of dopaminergic neurons (Figure 3C). Examination of these animals after exposure with IMI and FPF revealed that the number of mitochondrial puncta per CEP dendrite significantly increased (Figure 3D) while the average length of mitochondrial puncta significantly decreased (Figure 3E) compared to vehicle control, suggesting either a pesticide-induced increase in mitochondrial fission or a reduction in fusion. We also noticed that dendritic blebs occasionally coincided with mitochondrial puncta (see insets in Figure 3C), a phenomenon linked to neuronal excitotoxicity^58^.

**Figure 3.**
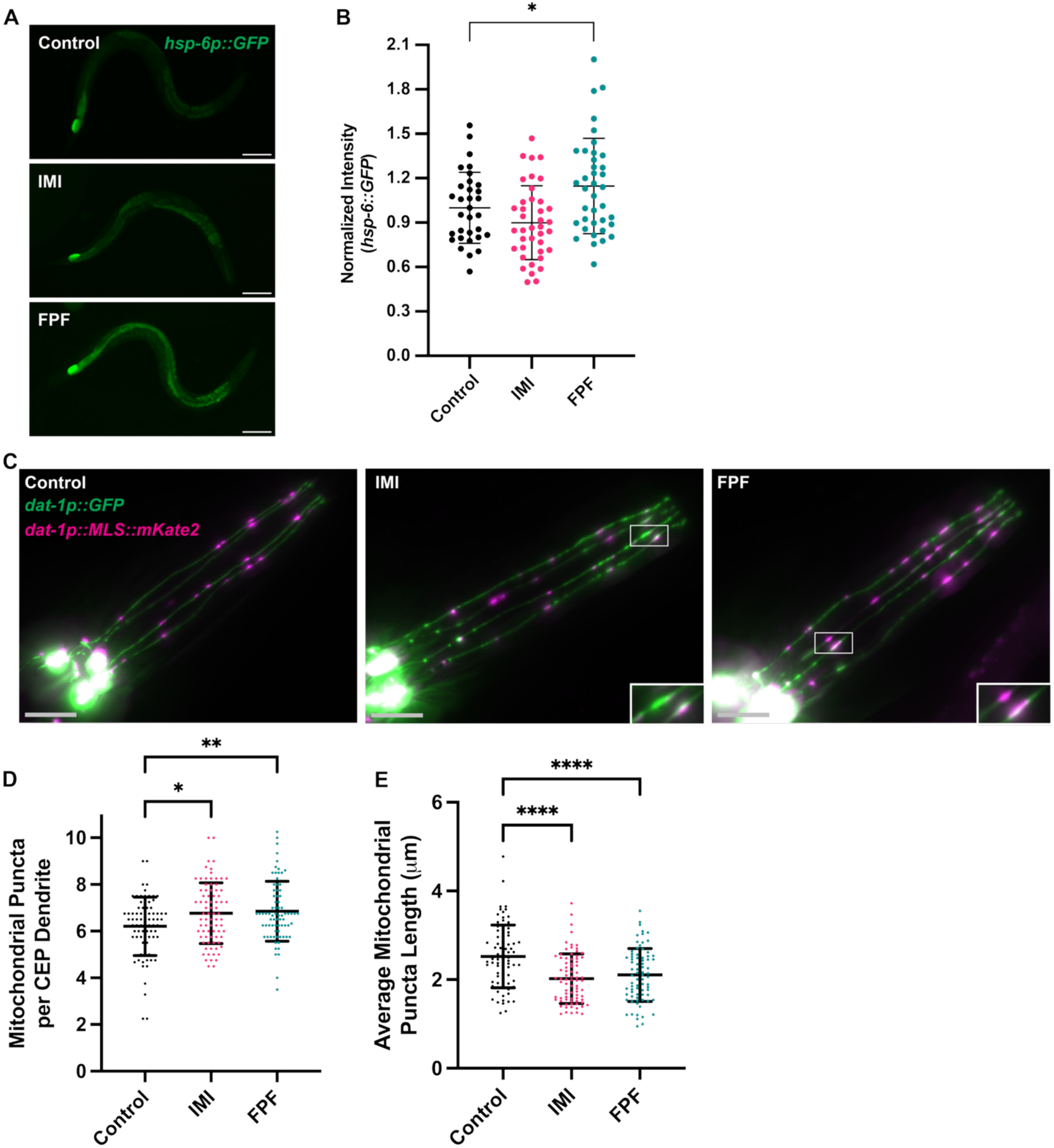
IMI and FPF elicit mitochondrial surveillance mechanisms. (A) Representative images of *hsp-6p::GFP* animals exposed to vehicle control (0.1% DMSO) or IMI/FPF (500μM) for 48 hours. Worms are oriented with the tail on the left, head on the right. Scale bar, 100μm. (B) Quantification of *hsp-6p::GFP* fluorescence intensity across each exposure condition. Corrected total worm fluorescence was used to determine intensity (see Methods for more details). One-way ANOVA with Dunnett’s multiple comparisons test was used to determine significance. * is p<0.05. Each dot is an individual animal. The line is the mean and error bars are standard deviations. N=2 exposures; n=32-38 animals. (C) Representative images of *dat1p::GFP* (green) + *dat-1p::MLS::mKate2* (magenta) animals exposed to vehicle control (0.1% DMSO) or IMI/FPF (500μM) for 48 hours. Four CEP neuronal bodies (bright white circles) and four CEP dendrites extending to the nose of the animal in the upper right are visible. Insets for IMI and FPF highlight blebs and mitochondrial puncta along CEP dendrites. Scale bar (lower left in grey), 20μm. (D) Quantification of the number of mitochondrial puncta per CEP dendrite across the exposure conditions. One-way ANOVA with Dunnett’s multiple comparisons test was used to determine significance. * is p<0.05 and ** is p<0.01. Each dot is an individual animal. The line is the mean and error bars are standard deviations. N=4 exposures; n=76-78 animals. (E) Quantification of the length of mitochondrial puncta on CEP dendrites across the exposure conditions. One-way ANOVA with Dunnett’s multiple comparisons test was used to determine significance. **** is p<0.0001. Each dot is an individual animal. The line is the mean and error bars are standard deviations. N=4 exposures; n=76-78 animals

Excess ROS can both result from, and cause, mitochondrial dysfunction, leading to oxidative stress^59^. Generation of ROS is common to many neurodegenerative diseases, including PD, and many pesticides trigger neuronal ROS^60^. In *C. elegans*, the transcription factor DAF-16 is known to translocate to the nucleus from the cytoplasm when *C. elegans* is exposed to ROS-generating conditions including starvation, heat, and paraquat^61^. Using a transgenic strain expressing a CRISPR allele of *daf-16* tagged at the C-terminus with GFP^62^, we found extensive nuclear translocation of DAF-16 in the head region of animals exposed to either IMI or FPF (Figures 4A and 4B). Previous studies have established that various ROS-generating chemicals cause a concomitant elevation in expression of the mitochondrial superoxide dismutase SOD-3 downstream of DAF-16^63^. Using a transgenic *C. elegans* strain carrying a *sod-3p::GFP* reporter, we found that both IMI and FPF cause a small, but significant increase in *sod-3* expression (Figures 4C and 4D). Increased expression was especially evident in the head (blue border insets Figure 4C) and tail (magenta border insets Figure 4C). To determine whether generation of ROS causes dopaminergic blebbing during IMI or FPF exposure, we used the potent antioxidant NAC, which alleviates ROS-mediated neuronal injury in a wide variety of contexts, including human dopaminergic neurons exposed to rotenone^64^. NAC administered at the same time as pesticide exposure was able to prevent the blebbiness induced by IMI (Figures 4E and 4F). NAC also counteracted FPF-induced blebbiness, though it remained elevated compared to control. Treatment with NAC also rescued the food race assay defect of IMI and FPF exposed animals (Figure 4G), suggesting that ROS generation is at least partially responsible for the neuronal blebbing and behavioral defects we observed.

Previous research in *C. elegans* determined that animals defective for DLK-1 pathway signaling via a loss-of-function mutation in the MAPKK MKK-4 have both reduced neurite regeneration, including CEP regeneration after laser surgery, and are more sensitive to 6-OHDA induced CEP neuron damage^40^. This is thought to occur via MKK-4 activation of PMK-3; however, MKK-4 can also activate the p38 MAPK PMK-1 in a DLK-1 independent manner^65^. The p38 pathway regulates oxidative stress responses, and inhibition of p38 exerts a protective effect in PD models^66,67^. To determine the role of MKK-4 in IMI and FPF-induced dopaminergic neurodegeneration, we exposed *mkk-4* loss-of-function mutants to IMI and FPF and measured CEP dendritic blebbiness and food race assay performance. Surprisingly, we found that *mkk-4* mutants showed remarkably lower blebbing after either pesticide exposure compared to control animals (Figures 4H and 4I). Additionally, *mkk-4* mutants displayed no foraging defects after IMI or FPF exposure compared to unexposed mutants (Figure 4J). These results therefore support the role of p38 MAPK in neurodegeneration and show that inhibition of a MAPKK upstream of p38 could be a potential therapeutic strategy for PD. Overall, this study establishes a mechanistic framework linking environmental nAChR agonist exposure to dopaminergic neurodegeneration, implicating MKK-4 mediated MAPK signaling and oxidative stress as key contributors.

**Figure 4.**
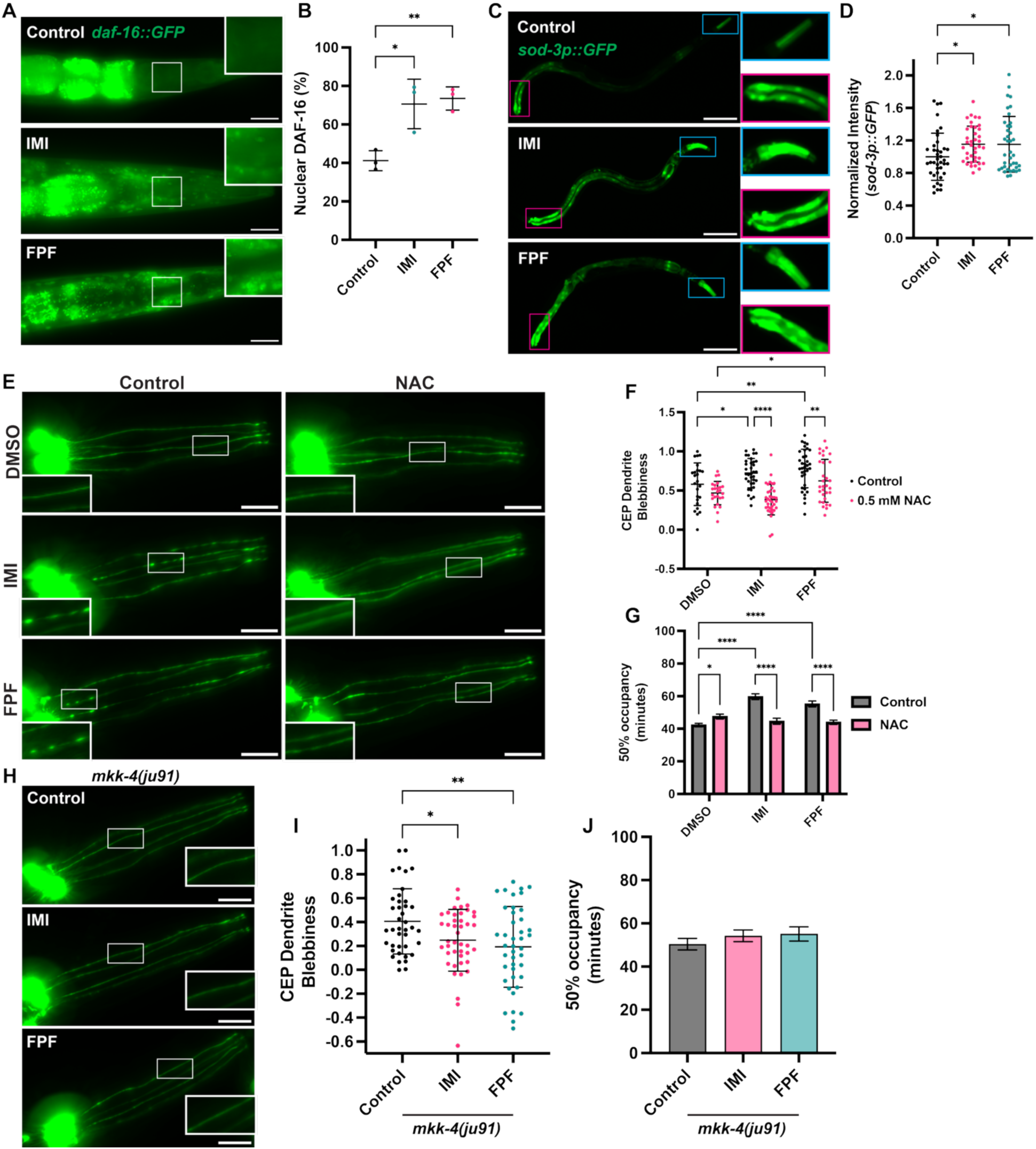
Antioxidant treatment and loss of *mkk-4* rescues IMI and FPF induced CEP dendritic blebbing. (A) Representative images of *daf-16::GFP* animals exposed to vehicle control (0.1% DMSO) or IMI/FPF (500μM) for 48 hours. Worms are oriented with nose tip on the right. Insets highlight increased expression in the nuclei in the head following IMI and FPF exposure. Scale bar, 30μm. (B) Quantification of DAF-16 nuclear translocation across each exposure condition. Percentages reflect the number of animals with nuclear DAF-16::GFP localization. One-way ANOVA with Dunnett’s multiple comparisons test was used to determine significance. * is p<0.05 and ** is p<0.01. Each dot is an exposure of at least 30 animals. The line is the mean and error bars are standard deviations. N=3 exposures. (C) Representative images of *sod-3p::GFP* animals exposed to vehicle control (0.1% DMSO) or IMI/FPF (500μM) for 48 hours. Worms are oriented with the tail on the left, head on the right. Blue-border insets highlight expression in the head, magenta-border insets highlight expression in the tail. Scale bar, 100μm. (D) Quantification of *sod-3p::GFP* fluorescence intensity across each exposure condition. Corrected total worm fluorescence was used to determine intensity (see Methods for more details). One-way ANOVA with Dunnett’s multiple comparisons test was used to determine significance. * is p<0.05. Each dot is an individual animal. The line is the mean and error bars are standard deviations. N=2 exposures; n=40 animals. (E) Representative images of *dat-1p::GFP* animals exposed to vehicle control (0.1% DMSO) or 500μM of IMI or FPF for 48 hours with (right column) or without (left column) NAC treatment. Scale bar, 20μm. (F) Quantification of CEP dendrite blebbiness of exposure groups shown in (E). See Methods section for details on blebbiness calculation. Two-way ANOVA with Tukey’s multiple comparisons test was used to determine significance. * is p<0.05, ** is p<0.01, and **** is p<0.0001. Each dot is an individual animal. The line is the mean and error bars are standard deviations. N=2 exposures; n=25-36 animals. (G) Using food race assays on the exposure groups shown in (E), the time it took half the animals to occupy the lawn was determined as in Figure 2F. The mean of each group is displayed. Error bars are standard deviation. Two-way ANOVA with Tukey’s multiple comparisons test was used to determine significance. * is p<0.05 and **** is p<0.0001. N=4 exposures. (H) Representative images of *mkk-4(ju91); dat-1p::GFP* animals exposed to vehicle control (0.1% DMSO) or 500μM of IMI or FPF for 48 hours Scale bar, 20μm. (I) Quantification of CEP dendrite blebbiness of exposure groups shown in (H). See Methods section for details on blebbiness calculation. One-way ANOVA with Dunnett’s multiple comparisons test was used to determine significance. * is p<0.05 and ** is p<0.01. Each dot is an individual animal. The line is the mean and error bars are standard deviations. N=2 exposures; n=40-43 animals. (J) Using food race assays on the exposure groups shown in (H), the time it took half the animals to occupy the lawn was determined as in (G). The mean of each group is displayed. Error bars are standard deviation. One-way ANOVA with Dunnett’s multiple comparisons test was used to determine significance. N=4 exposures.

While we think it is likely that MKK-4 is exerting an effect via activation of PMK-1, we cannot rule out a role for PMK-3. Additionally, the role of other PMK-1 MAPKKs, namely SEK-1, in our model is unknown. Future experiments will be required to determine the role of these MAPK signaling pathways in more detail. Interestingly, a previous study reported that rotenone activates p38-mediated innate immune responses in intestinal cells of *C. elegans*, and that enhancing these pathways suppresses rotenone-induced dopaminergic neuron loss^68^. Conversely, loss of function mutations in *tir-1* (the *C. elegans* homolog of SARM1) and the associated downstream kinases *nsy-1, sek-1,* and *pmk-1* suppress motor neuron degeneration, pointing to the complex relationship of p38 MAPK signaling pathways and neurodegeneration^69^. Previous PD studies have shown promising results by inhibiting p38 either directly via blocking its kinase activity^67^ or indirectly by inhibiting upstream MAPKKs^66^. Ongoing clinical trials of the direct p38α inhibitor VX-745 (neflamapimod) have also shown some positive results for the closely related disease dementia with Lewy bodies (DLB)^70^. It is therefore likely that the level of p38 MAPK activity represents a balance between neuroprotection and neurodegeneration^71^. Tuning this activity after exposure to environmental stressors such as IMI or FPF exposure could be a potential therapeutic avenue.

The involvement of MAPK signaling pathways in PD and in our model of IMI and FPF-induced dopaminergic neurodegeneration is likely due to oxidative stress. FPF, though not IMI, caused an elevation in expression of the UPRmt-associated gene *hsp-6* (Figures 3A and 3B). Both pesticides increased CEP dendritic mitochondria numbers while decreasing their size (Figures 3C-E), indicative of defects in mitochondrial fission, fusion or both, while also causing nuclear translocation of *daf-16* and increased expression of its downstream target, the superoxide dismutase *sod-3* (Figures 4A-C). Most importantly, co-administering the antioxidant NAC during IMI or FPF exposure alleviated both CEP dendritic blebbing and foraging defects (Figures 4D-F). Antioxidant administration in preclinical PD models is often successful, though clinical trials have shown mixed and sometimes poor results^72^. These clinical failures are most likely due to a combination of factors that include timing of the treatment in relation to disease progression, short antioxidant half-lives, low bioavailability, limited blood-brain barrier penetration, and lack of targeted delivery to affected neurons. Our data indicates that, although the oxidative stress response is systemic, IMI and FPF elicits blebbing only in dopaminergic neurons. This is in line with studies showing that dopaminergic neurons are sensitive to oxidative stress due to a combination of α-synuclein aggregation, increased energetic demand stemming from extensive axonal arborization, calcium-related pacemaking, and oxidation or metabolism of cytosolic dopamine^73^. *C. elegans* lacks an α-synuclein homolog and its dopaminergic neurons do not have extensive arborization or pacemaking activity, suggesting that oxidation or metabolism of cytosolic dopamine could be driving their susceptibility to IMI and FPF. It will be interesting to determine if IMI and FPF, which likely bind to nAChRs on the CEP neurons causing a rise in calcium influx, also results in cytosolic dopamine accumulation, furthering their excitotoxicity. The wide variety of genetic tools available in *C. elegans* will also allow the investigation of how IMI and FPF exposure interacts with known PD genes, many of which are involved in mitochondrial function, oxidative stress response, and kinase activity^17–20^.

## Methods

### Key resources table

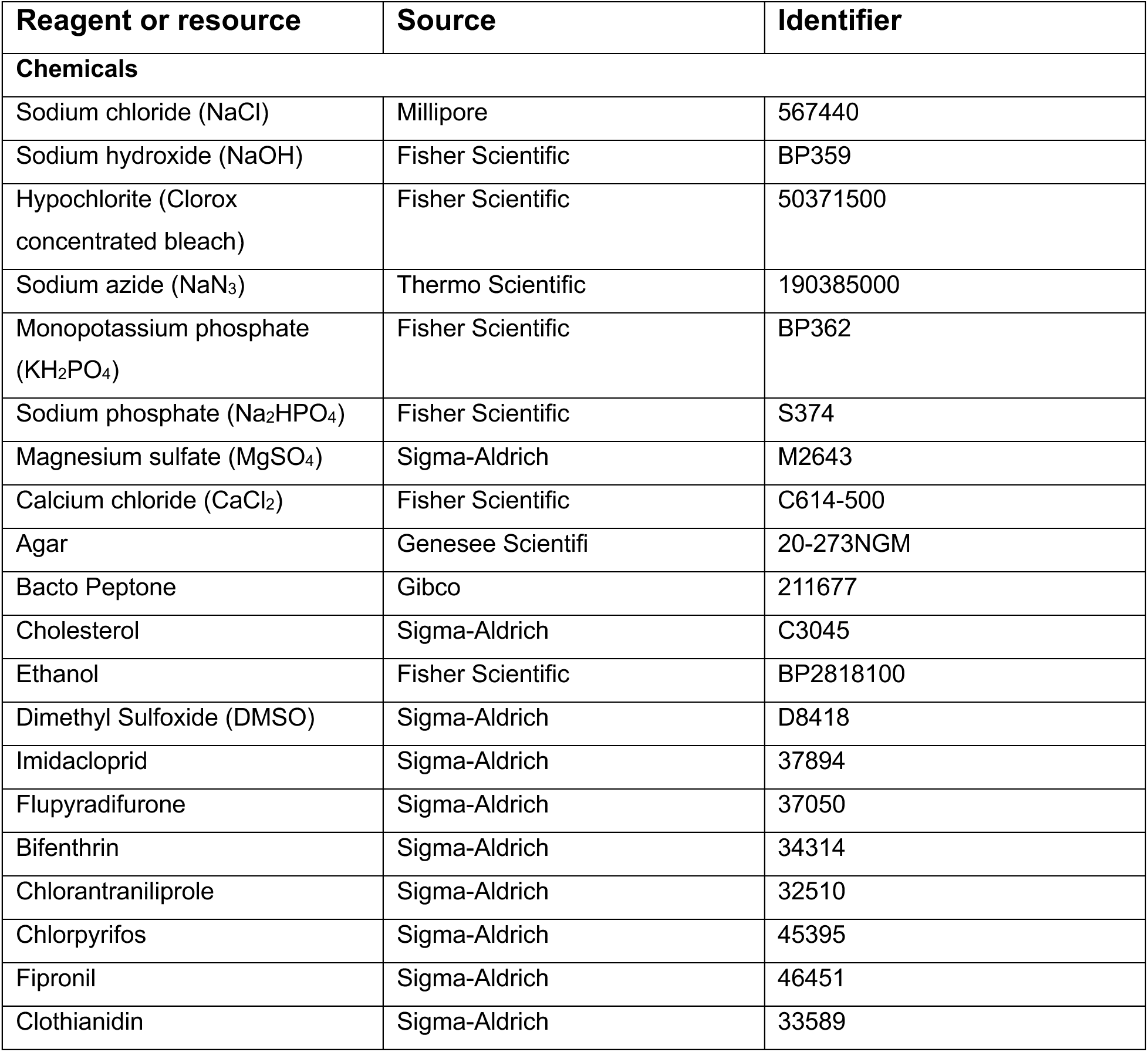
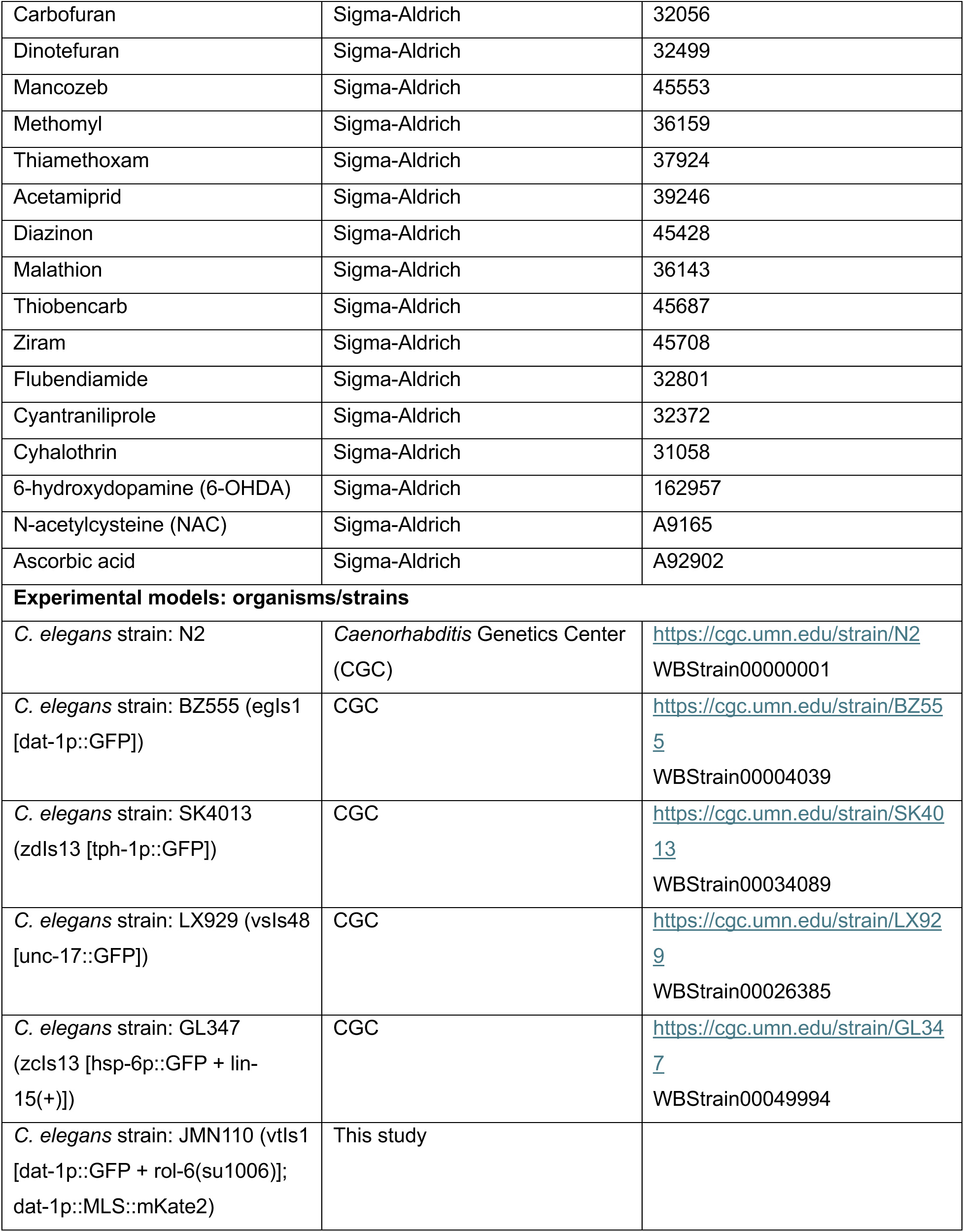
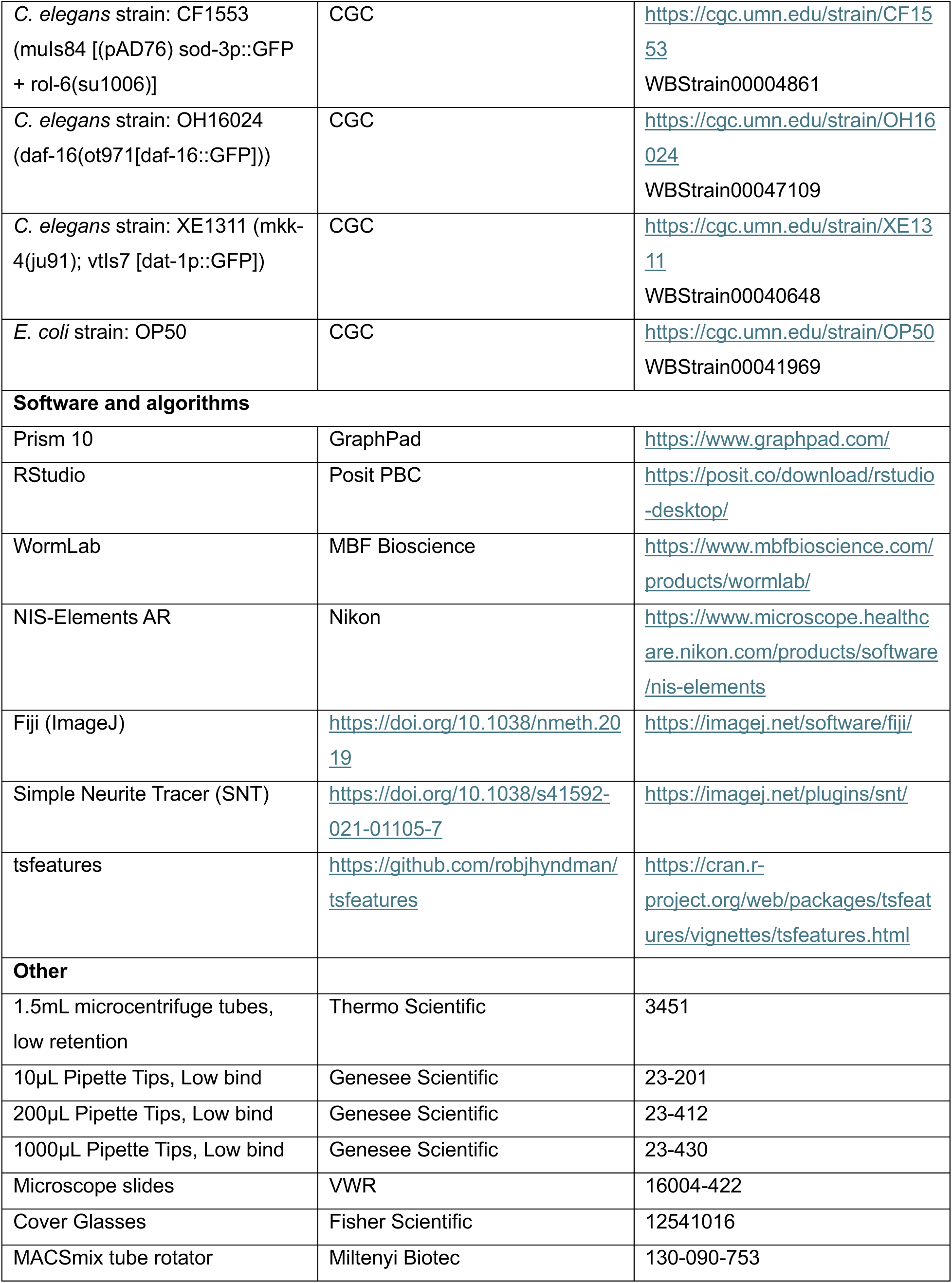
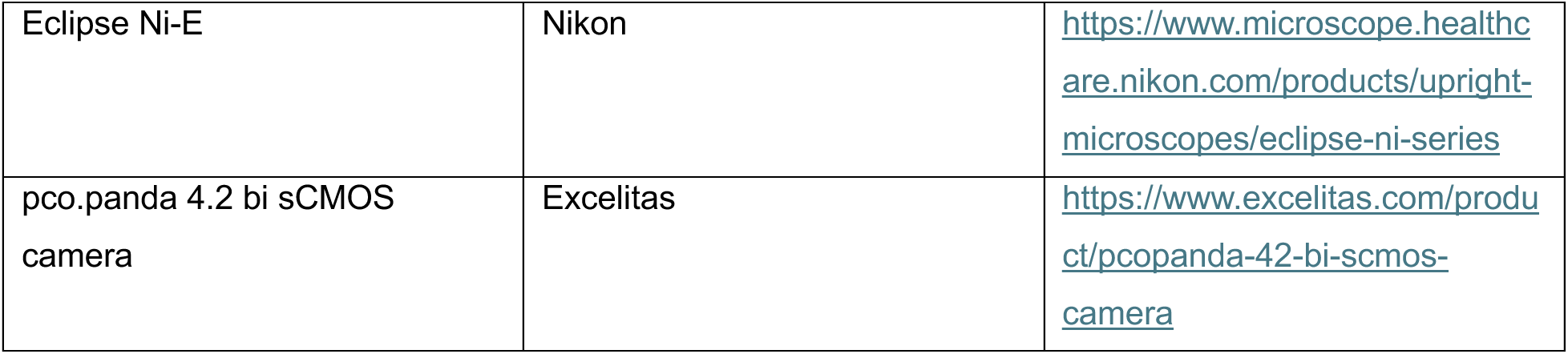

### Resource availability

#### Lead contact

Further information and requests for resources and reagents should be directed to and will be fulfilled by the lead contact, Dr. Patrick Allard.

#### Materials availability

All plasmids, strains, and other reagents generated in this study are freely available upon request.

#### Data and code availability

Data are reported in the figures of the paper and supplemental information. No original code was reported in the paper. Any additional information required to reanalyze the data reported in this paper is available from the lead contact upon request.

### Experimental model and subject details

N2 (wild type) worms were used for lethality and behavioral experiments. The strains BZ555 *dat-1p:GFP,* SK4013 *tph-1p::GFP,* and LX929 *unc-17p::GFP* were used for dopaminergic, serotoninergic, and cholinergic neuron visualization, respectively. Strain GL347 *hsp-6p::GFP* was used to visualize *hsp-6* expression. For dopaminergic mitochondria experiments the strain JMN110 vtIs1 [*dat-1p::GFP + rol-6(su1006)*] *+ dat-1p::MLS::mKate2* was used. The strains CF1553 *sod-3p::GFP + rol-6(su1006)* and OH16024 *daf-16(ot971[daf-16::GFP])* were used to visualize *sod-3* expression and *daf-16* translocation, respectively. For *mkk-4* experiments, strain XE1311 *mkk-4(ju91) + dat-1::GFP* was used. Worms were cultured on standard nematode growth medium (NGM) plates seeded with single colony OP50 *E. coli* and maintained at 20°C.

### Method details

#### C. *elegans* pesticide exposure

For pesticide exposures, a population of gravid adult worms was bleached. The embryos obtained were plated on standard OP50 seeded NGM plates and allowed to grow to the L4 stage (approximately 50 hours post bleaching). Nematodes were exposed for 48 hours in liquid culture containing M9 buffer solution, standard OP50 bacteria (10mg/mL), and pesticides diluted in DMSO (500mM stock solutions) at final concentrations of 500μM, 166μM, 55μM, or 18μM in 1.5mL microcentrifuge tubes. 6-OHDA (100mM stock solution in water) exposures were done at a final concentration of 10mM with 4mM ascorbic acid to prevent auto-oxidation of 6-OHDA. For NAC experiments, NAC was added along with pesticides for a final NAC concentration of 0.5mM. Tubes were placed in a MACSmix Tube Rotator (Miltenyi Biotec 130-090-753) in a 20°C incubator. After exposure tubes were washed three times with M9 buffer, with animals settling to the bottom by gravity in between washes, before downstream experiments were conducted.

#### Survival assays

Exposed and washed worms were transferred to NGM plates without OP50 and immediately counted for survival. Survival index was calculated as (total number of animals – animals that did not respond to a gentle touch with a sterilized platinum wire) / total number of animals.

#### Neurodegeneration imaging and quantification

Exposed and washed worms were transferred to microscopy slides, anesthetized with 5μL 0.5M sodium azide, and imaged immediately. Images of CEP dendrites were collected at 0.8μm z-intervals with an Eclipse Ni-E (Nikon) equipped with a 40x objective and a pco.panda 4.2 bi sCMOS camera (Excelitas) controlled by the NIS-Elements AR software (Nikon). Captured images were analyzed using ImageJ software. Maximum intensity projections of z-stack images were made, converted into 8-bit grayscale, and then opened in the Simple Neurite Tracer (SNT) plugin of ImageJ. Using SNT, traces were made along the length of each CEP dendrite starting at the neuronal cell body and ending at the tip near the nose of the animal. Smoothened fluorescence intensity profiles were generated for each dendrite from these traces using a solid disk with radius of 3 pixels for the sampling neighborhood and mean fluorescence intensity for the integration metric. Blebs along the dendrites showed up as spikes in the fluorescence intensity profiles and so a seasonal-trend decomposition using LOESS (STL) method was adapted to analyze the spikiness. The stl_features function in the tsfeatures package in R was used to compute the spike (variance of the leave-one-out variances of the remainder component of the decomposition) of each dendrite’s fluorescence intensity profile. Spike measurements of dendrites were then averaged for each animal, log10 normalized, and min-max scaled to 0-1 relative to control to obtain CEP dendrite blebbiness. Images of CEP neurons, along with serotoninergic HSN neurons and cholinergic ventral motor neurons, were also subjected to blind scoring. For blind scoring, two independent scorers blinded to the origin of images determined the level of blebbing using the following definitions: None: no blebs observed, Mild: at least one bleb on one dendrite, Moderate: multiple blebs on at least one dendrite, Severe: multiple blebs on multiple dendrites.

#### Basal slowing and foraging

To measure basal slowing, exposed animals were allowed to recover on NGM plates with OP50 *E. coli* for 10 minutes. Half of the animals were then transferred to NGM plates without OP50 (No Food) and half to NGM plates with OP50 (Food) to create paired trials. After 10 minutes of acclimatization, 1-minute videos of animals moving on the plates were recorded using the WormLab imaging system (MBF Bioscience). Velocities of individual worms were measured using WormLab software. Crawling speed was calculated as the average velocity for each exposure, and basal slowing speed was calculated as the difference in crawling speed between the paired no food and food assays. To measure foraging, a food race assay was used. Animals were allowed to recover on NGM plates with OP50 for 10 minutes and then transferred to the side of an NGM plate opposite a 20μL lawn of OP50. The percentage of animals reaching the food was calculated as: number of animals on the lawn of OP50 / total number of animals. This was measured every 10 minutes for 2 hours. The 50% occupancy time was determined from these measurements using a variable slope model.

#### Quantitative fluorescence measurements

For *hsp-6p::GFP* and *sod-3p::GFP* measurements, images were loaded into ImageJ and regions of interest were drawn around whole worms. Area and integrated density were measured for each worm. The average mean fluorescence intensity of three random background regions was also calculated for each exposure. To account for possible differences in worm size, the fluorescence intensity of each worm was measured as the corrected total worm fluorescence (CTWF), where CTWF = Integrated Density – (Area of selected worm x average mean fluorescence of background readings). CTWF measurements were normalized to the control.

#### C. *elegans* strain generation

To generate the *dat-1p::MLS::mKate2* plasmid, mKate2 was amplified from a previously *C. elegans* optimized plasmid. The mitochondrial localization sequence (MLS) was originally amplified from Fire Vector pPD96.32. pPD96.32 was a gift from Andrew Fire (Addgene plasmid # 1504; http://n2t.net/addgene:1504; RRID:Addgene_1504). The amplified mKate2 sequence was inserted into the plasmid containing the *dat-1* promoter and the MLS, and insertion was confirmed by colony PCR. The plasmid was sequenced to check for mutations and co-injected with 50ng/μL *unc-119* rescue DNA, 50ng/μL pBsSK, and 50ng/μL EcoR1 cut salmon sperm DNA into *unc-119(ed4)* hermaphrodites. Once extrachromosomal lines were established and *dat-1p::MLS::mKate2* signal was confirmed, the plasmid was integrated by gamma irradiation as previously described. Integrated lines were outcrossed with N2 6x to remove possible background mutations. Subsequently, *dat-1p::MLS::mKate2* was crossed with BY200 (vtIs1 [*dat-1p::GFP + rol-6(su1006)]* by standard mating practices to generate JMN110.

#### Mitochondria quantification

To measure the number and size of mitochondria in CEP dendrites, strain JMN110 vtIs1 [*dat-1p::GFP + rol-6(su1006)*] *+ dat-1p::MLS::mKate2* was used. Images were loaded into ImageJ and straight lines were drawn through mitochondria puncta along CEP dendrites to obtain length. Puncta were counted using the Cell Counter plugin.

## Acknowledgements

This work was funded by NIH grants T32ES015457 to A.R.F and P.A, T32ES021432 to K.S.M, P42ES010356 and R01ES034270 to J.N.M, and R35GM118049 to D.R.S.

## Supplemental Figures

**Figure S1.**
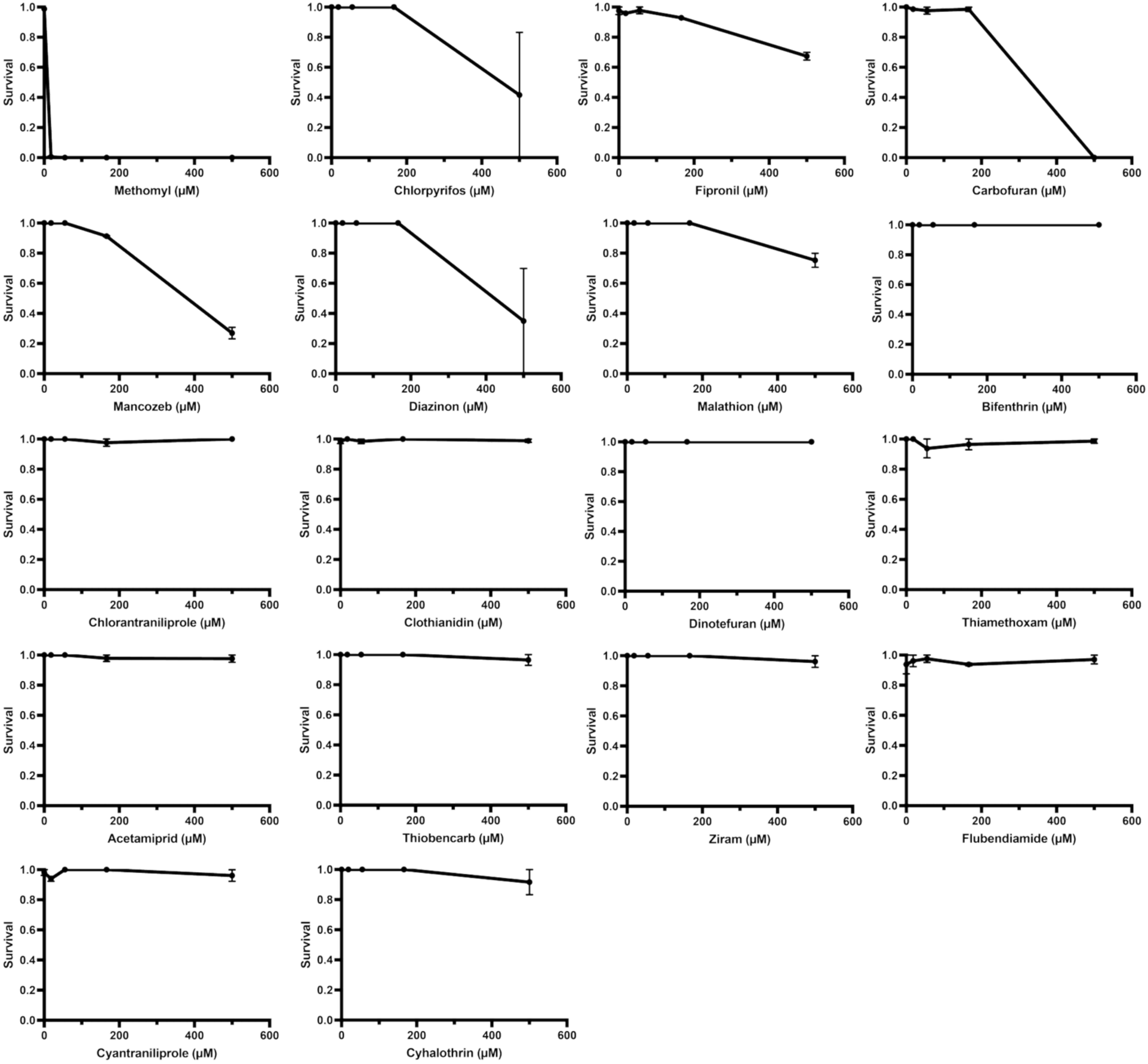
Survival assays for 18 different pesticides. Animals were exposed as in Figure 1 to 18 different pesticides and survival was checked immediately after exposure. Dots are the mean at each concentration and error bars are SEM. N=2 exposures.

**Figure S2.**
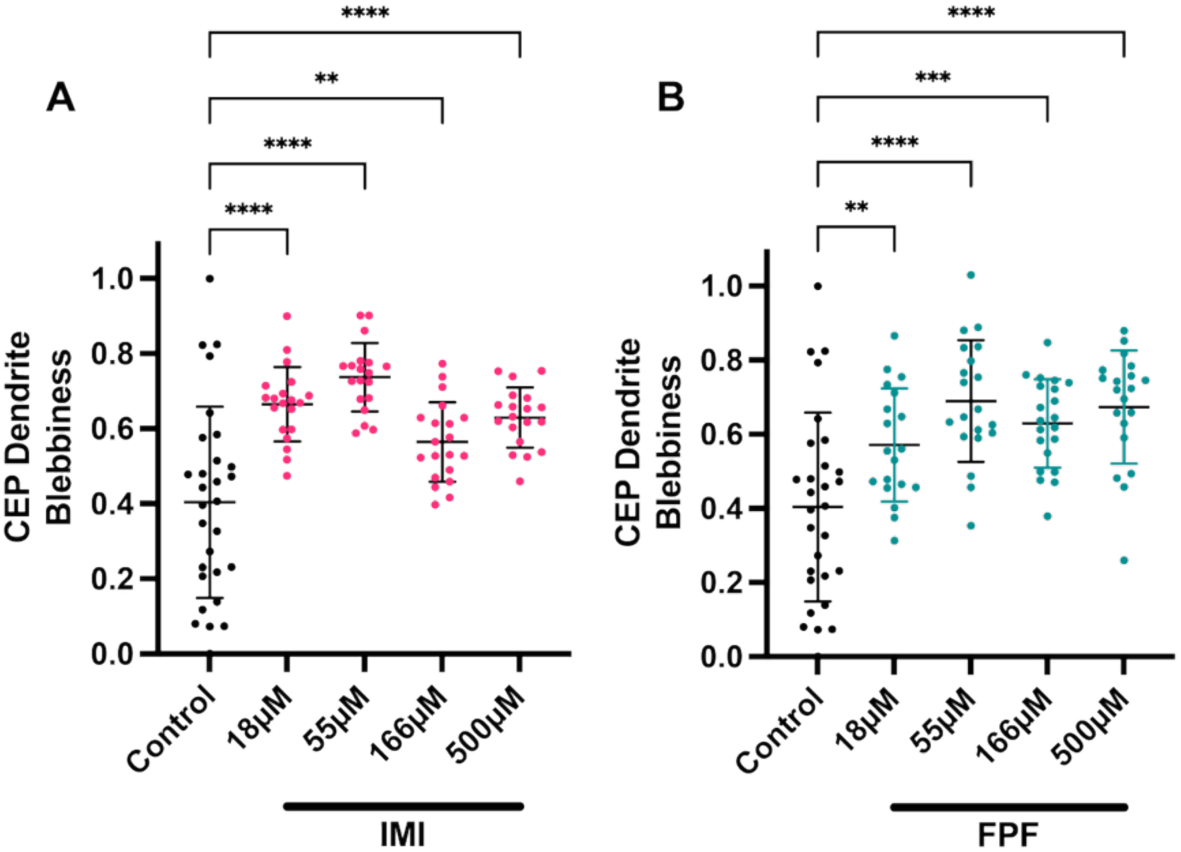
IMI and FPF cause CEP dendrite blebbing from 18-500μM. (A and B) Animals were exposed for 48 hours as in Figure 1 to vehicle control (0.1% DMSO), IMI (A), or FPF (B) at the indicated concentrations. Quantification of blebbiness of CEP dendrites after exposure was performed as in Figure 1. One-way ANOVA with Dunnett’s multiple comparisons test was used to determine significance. ** is p<0.01, *** is p<0.001, and **** is p<0.0001. Each dot is an individual animal. The line is the mean and error bars are standard deviations. N=2-3 exposures; n=19-29 animals.

**Figure S3.**
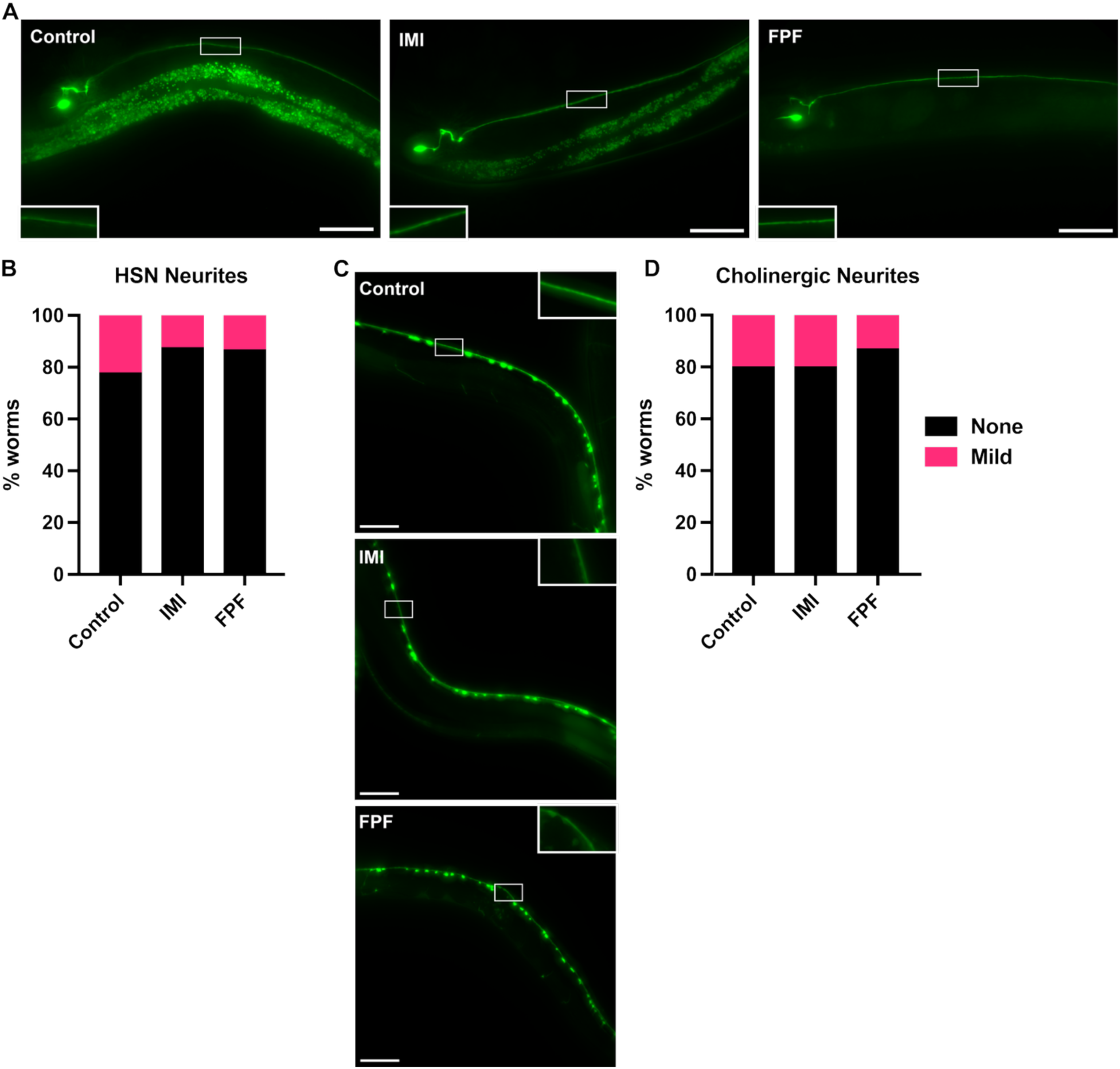
IMI and FPF do not cause blebbing in serotonergic or cholinergic neurons. (A) Representative images of serotonergic HSN neurons in *tph-1::GFP* animals after 48-hour exposure to vehicle control (0.1% DMSO) or IMI/FPF (500μM). HSN cell body is visible on the left, with an axonal projection around the vulva and along the ventral nerve cord moving towards the head on the right. Insets highlight HSN neurites. Scale bar, 50μm (B) Blind scoring of HSN neurites after 48-hour exposure to vehicle control (0.1% DMSO) or IMI/FPF (500μM). Two blinded scorers determined the level of blebbing using the following definitions: None: no blebs observed, Mild: at least one bleb on one dendrite, Moderate: multiple blebs on at least one dendrite, Severe: multiple blebs on multiple dendrites. N=2 exposures; n=20-30 animals. (C) Representative images of cholinergic motor neurons in *unc-17p::GFP* animals after exposure as in (A). Ventral nerve cord neuronal cell bodies and projections (highlighted in insets) are visible. Scale bar, 50μm. (D) Blind scoring of cholinergic neurites as in (B). N=2 exposures, n =16-30 animals.

**Figure S4.**
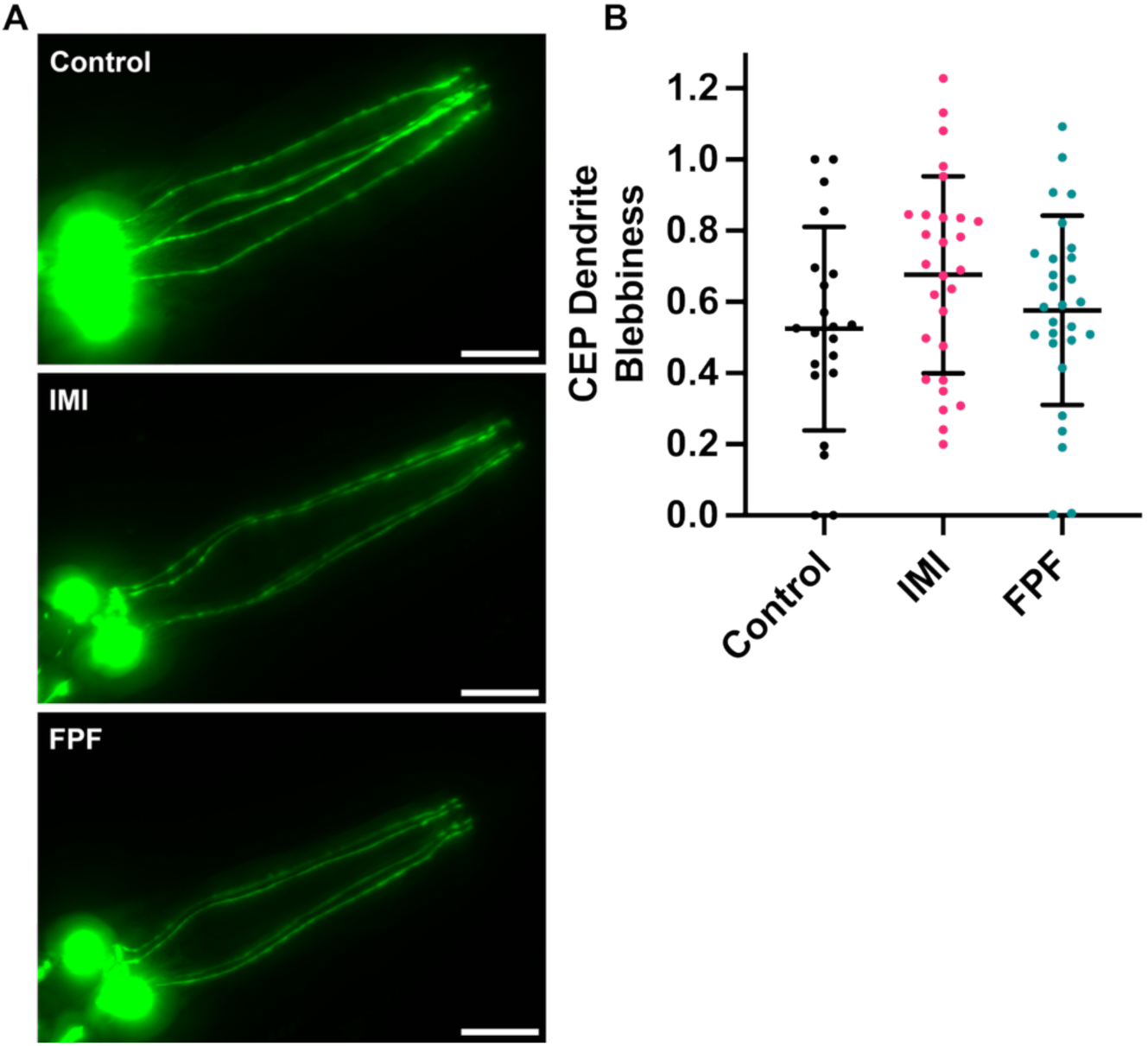
L1-L4 exposure to IMI or FPF does not cause CEP dendritic blebbing. (A) Representative images of dopaminergic CEP dendrites in *dat-1p::GFP* animals after 48-hour exposure to vehicle control (0.1% DMSO) or IMI/FPF (500μM) from larval stage L1 to L4. Scale bar, 20μm. Quantification of blebbiness of CEP dendrites after exposure was performed as in Figure 1. One-way ANOVA with Dunnett’s multiple comparisons test was used to determine significance. Each dot is an individual animal. The line is the mean and error bars are standard deviations. N=2 exposures; n=21-28 animals.

## References

1. Brown, T.P., Rumsby, P.C., Capleton, A.C., Rushton, L., and Levy, L.S. (2006). Pesticides and Parkinson’s disease--is there a link? Environ Health Perspect 114, 156–164. 10.1289/ehp.8095.

2. Wang, A., Costello, S., Cockburn, M., Zhang, X., Bronstein, J., and Ritz, B. (2011). Parkinson’s disease risk from ambient exposure to pesticides. Eur J Epidemiol 26, 547– 555. 10.1007/s10654-011-9574-5.

3. Kamel, F., Tanner, C., Umbach, D., Hoppin, J., Alavanja, M., Blair, A., Comyns, K., Goldman, S., Korell, M., Langston, J., et al. (2007). Pesticide Exposure and Self-reported Parkinson’s Disease in the Agricultural Health Study. Am J Epidemiol 165, 364–374. 10.1093/aje/kwk024.

4. Arellano, M., Barnhill, L.M., Kim, A.M., Mahmudul Hasan, K.M., Li, S., Paul, K.C., Peng, C., Ritz, B., and Bronstein, J.M. (2026). Identification of pesticides associated with an increased risk of Parkinson’s disease using a multi-screen approach. Environment International 208, 110087. 10.1016/j.envint.2026.110087.

5. Paul, K.C., and Ritz, B. (2022). Epidemiology meets toxicogenomics: Mining toxicologic evidence in support of an untargeted analysis of pesticides exposure and Parkinson’s disease. Environ Int 170, 107613. 10.1016/j.envint.2022.107613.

6. Costas-Ferreira, C., and Faro, L.R.F. (2021). Neurotoxic Effects of Neonicotinoids on Mammals: What Is There beyond the Activation of Nicotinic Acetylcholine Receptors?—A Systematic Review. Int J Mol Sci 22, 8413. 10.3390/ijms22168413.

7. Loser, D., Hinojosa, M.G., Blum, J., Schaefer, J., Brüll, M., Johansson, Y., Suciu, I., Grillberger, K., Danker, T., Möller, C., et al. (2021). Functional alterations by a subgroup of neonicotinoid pesticides in human dopaminergic neurons. Arch Toxicol 95, 2081–2107. 10.1007/s00204-021-03031-1.

8. Sinclair, P., Hakeem, J., Kumar, S.G., Loser, D., Dixit, K., Leist, M., Kraushaar, U., and Kabbani, N. (2023). Proteomic responses in the human dopaminergic LUHMES cell line to imidacloprid and its metabolites imidacloprid-olefin and desnitro-imidacloprid. Pesticide Biochemistry and Physiology 194, 105473. 10.1016/j.pestbp.2023.105473.

9. Nauen, R., Jeschke, P., Velten, R., Beck, M.E., Ebbinghaus-Kintscher, U., Thielert, W., Wölfel, K., Haas, M., Kunz, K., and Raupach, G. (2015). Flupyradifurone: a brief profile of a new butenolide insecticide. Pest Manag Sci 71, 850–862. 10.1002/ps.3932.

10. Kathage, J., Castañera, P., Alonso-Prados, J.L., Gómez-Barbero, M., and Rodríguez-Cerezo, E. (2018). The impact of restrictions on neonicotinoid and fipronil insecticides on pest management in maize, oilseed rape and sunflower in eight European Union regions. Pest Manag Sci 74, 88–99. 10.1002/ps.4715.

11. Hogberg, H.T., Frische, E., Gröters, S., Kovi, R.C., Rao, D.B., Terron, A., and Winter, M.J. (2025). 2024 International Academy of Toxicologic Pathology (IATP) Satellite Symposium: New Approach Methodologies (NAMs) for Neurotoxicity Assessment and Regulatory Perspectives. Toxicol Pathol 53, 305–320. 10.1177/01926233251335719.

12. Hunt, P.R. (2017). The C. elegans model in toxicity testing. J Appl Toxicol 37, 50–59. 10.1002/jat.3357.

13. Cooper, J.F., and Van Raamsdonk, J.M. Modeling Parkinson’s Disease in C. elegans. J Parkinsons Dis 8, 17–32. 10.3233/JPD-171258.

14. Hunt, P.R., Camacho, J.A., and Sprando, R.L. (2020). *Caenorhabditis elegans* for predictive toxicology. Current Opinion in Toxicology 23–24, 23–28. 10.1016/j.cotox.2020.02.004.

15. Willis, A.W. (2013). Parkinson Disease in the Elderly Adult. Mo Med 110, 406–410.

16. Surmeier, D.J., Obeso, J.A., and Halliday, G.M. (2017). Selective neuronal vulnerability in Parkinson disease. Nat Rev Neurosci 18, 101–113. 10.1038/nrn.2016.178.

17. Beilina, A., and Cookson, M.R. (2016). Genes associated with Parkinson’s disease: regulation of autophagy and beyond. J Neurochem 139, 91–107. 10.1111/jnc.13266.

18. Kumaran, R., and Cookson, M.R. (2015). Pathways to Parkinsonism Redux: convergent pathobiological mechanisms in genetics of Parkinson’s disease. Hum Mol Genet 24, R32–R44. 10.1093/hmg/ddv236.

19. Lin, M.K., and Farrer, M.J. (2014). Genetics and genomics of Parkinson’s disease. Genome Med 6, 48. 10.1186/gm566.

20. Mullin, S., and Schapira, A. (2015). The genetics of Parkinson’s disease. Br Med Bull 114, 39–52. 10.1093/bmb/ldv022.

21. Bogers, J.S., Bloem, B.R., and Den Heijer, J.M. The Etiology of Parkinson’s Disease: New Perspectives from Gene-Environment Interactions. J Parkinsons Dis 13, 1281–1288. 10.3233/JPD-230250.

22. Berry, C., La Vecchia, C., and Nicotera, P. (2010). Paraquat and Parkinson’s disease. Cell Death Differ 17, 1115–1125. 10.1038/cdd.2009.217.

23. Xiong, N., Long, X., Xiong, J., Jia, M., Chen, C., Huang, J., Ghoorah, D., Kong, X., Lin, Z., and Wang, T. (2012). Mitochondrial complex I inhibitor rotenone-induced toxicity and its potential mechanisms in Parkinson’s disease models. Crit Rev Toxicol 42, 613–632. 10.3109/10408444.2012.680431.

24. Cristóvão, A.C., Campos, F.L., Je, G., Esteves, M., Guhathakurta, S., Yang, L., Beal, M.F., Fonseca, B.M., Salgado, A.J., Queiroz, J., et al. (2020). Characterization of a Parkinson’s disease rat model using an upgraded paraquat exposure paradigm. Eur J Neurosci 52, 3242–3255. 10.1111/ejn.14683.

25. Cannon, J.R., Tapias, V.M., Na, H.M., Honick, A.S., Drolet, R.E., and Greenamyre, J.T. (2009). A highly reproducible rotenone model of Parkinson’s disease. Neurobiol Dis 34, 279–290. 10.1016/j.nbd.2009.01.016.

26. Van Laar, A.D., Webb, K.R., Keeney, M.T., Van Laar, V.S., Zharikov, A., Burton, E.A., Hastings, T.G., Glajch, K.E., Hirst, W.D., Greenamyre, J.T., et al. (2023). Transient exposure to rotenone causes degeneration and progressive parkinsonian motor deficits, neuroinflammation, and synucleinopathy. npj Parkinsons Dis. 9, 121. 10.1038/s41531-023-00561-6.

27. Godfray, H.C.J., Blacquière, T., Field, L.M., Hails, R.S., Petrokofsky, G., Potts, S.G., Raine, N.E., Vanbergen, A.J., and McLean, A.R. (2014). A restatement of the natural science evidence base concerning neonicotinoid insecticides and insect pollinators. Proc Biol Sci 281, 20140558. 10.1098/rspb.2014.0558.

28. Gross, M. (2014). Systemic pesticide concerns extend beyond the bees. Current Biology 24, R717–R720. 10.1016/j.cub.2014.07.071.

29. Manjon, C., Troczka, B.J., Zaworra, M., Beadle, K., Randall, E., Hertlein, G., Singh, K.S., Zimmer, C.T., Homem, R.A., Lueke, B., et al. (2018). Unravelling the Molecular Determinants of Bee Sensitivity to Neonicotinoid Insecticides. Current Biology 28, 1137–1143.e5. 10.1016/j.cub.2018.02.045.

30. Thompson, D.A., Lehmler, H.-J., Kolpin, D.W., Hladik, M.L., Vargo, J.D., Schilling, K.E., LeFevre, G.H., Peeples, T.L., Poch, M.C., LaDuca, L.E., et al. (2020). A critical review on the potential impacts of neonicotinoid insecticide use: current knowledge of environmental fate, toxicity, and implications for human health. Environ Sci Process Impacts 22, 1315–1346. 10.1039/c9em00586b.

31. Ma, Y., and Wang, Q. (2025). Neonicotinoid-Induced Cytotoxicity: Insights into Cellular Mechanisms and Health Risks. Toxics 13, 576. 10.3390/toxics13070576.

32. Tosi, S., and Nieh, J.C. (2019). Lethal and sublethal synergistic effects of a new systemic pesticide, flupyradifurone (Sivanto®), on honeybees. Proc Biol Sci 286, 20190433. 10.1098/rspb.2019.0433.

33. Hsiung, C., Boyd, P., Lemieux, H., Maisonneuve, F., Scroggins, R., and Princz, J. (2026). Fate and effects of flupyradifurone and sulfoxaflor on non-target soil-dwelling invertebrates. Environ Toxicol Chem, vgag053. 10.1093/etojnl/vgag053.

34. Siviter, H., and Muth, F. (2020). Do novel insecticides pose a threat to beneficial insects? Proc Biol Sci 287, 20201265. 10.1098/rspb.2020.1265.

35. Fang, N., Lu, Z., Hou, Z., Zhang, C., and Zhao, X. (2022). Hydrolysis and photolysis of flupyradifurone in aqueous solution and natural water: Degradation kinetics and pathway. Chemosphere 298, 134294. 10.1016/j.chemosphere.2022.134294.

36. Nass, R., Hall, D.H., Miller, D.M., and Blakely, R.D. (2002). Neurotoxin-induced degeneration of dopamine neurons in Caenorhabditis elegans. Proceedings of the National Academy of Sciences 99, 3264–3269. 10.1073/pnas.042497999.

37. Pu, P., and Le, W. (2008). Dopamine neuron degeneration induced by MPP+ is independent of CED-4 pathway in Caenorhabditis elegans. Cell Res 18, 978–981. 10.1038/cr.2008.279.

38. Masoudi, N., Ibanez-Cruceyra, P., Offenburger, S.-L., Holmes, A., and Gartner, A. (2014). Tetraspanin (TSP-17) Protects Dopaminergic Neurons against 6-OHDA-Induced Neurodegeneration in C. elegans. PLOS Genetics 10, e1004767. 10.1371/journal.pgen.1004767.

39. Zhou, S., Wang, Z., and Klaunig, J.E. (2013). Caenorhabditis elegans neuron degeneration and mitochondrial suppression caused by selected environmental chemicals. Int J Biochem Mol Biol 4, 191–200.

40. González-Hunt, C.P., Leung, M.C.K., Bodhicharla, R.K., McKeever, M.G., Arrant, A.E., Margillo, K.M., Ryde, I.T., Cyr, D.D., Kosmaczewski, S.G., Hammarlund, M., et al. (2014). Exposure to Mitochondrial Genotoxins and Dopaminergic Neurodegeneration in Caenorhabditis elegans. PLoS One 9, e114459. 10.1371/journal.pone.0114459.

41. Braungart, E., Gerlach, M., Riederer, P., Baumeister, R., and Hoener, M.C. (2004). Caenorhabditis elegans MPP+ model of Parkinson’s disease for high-throughput drug screenings. Neurodegener Dis 1, 175–183. 10.1159/000080983.

42. Zhou, S., Wang, Z., and Klaunig, J.E. (2013). Caenorhabditis elegans neuron degeneration and mitochondrial suppression caused by selected environmental chemicals. Int J Biochem Mol Biol 4, 191–200.

43. Sawin, E.R., Ranganathan, R., and Horvitz, H.R. (2000). C. elegans Locomotory Rate Is Modulated by the Environment through a Dopaminergic Pathway and by Experience through a Serotonergic Pathway. Neuron 26, 619–631. 10.1016/S0896-6273(00)81199-X.

44. Hills, T., Brockie, P.J., and Maricq, A.V. (2004). Dopamine and Glutamate Control Area-Restricted Search Behavior in Caenorhabditis elegans. J Neurosci 24, 1217–1225. 10.1523/JNEUROSCI.1569-03.2004.

45. Lakso, M., Vartiainen, S., Moilanen, A.-M., Sirviö, J., Thomas, J.H., Nass, R., Blakely, R.D., and Wong, G. (2003). Dopaminergic neuronal loss and motor deficits in Caenorhabditis elegans overexpressing human α-synuclein. Journal of Neurochemistry 86, 165–172. 10.1046/j.1471-4159.2003.01809.x.

46. Yao, C., El Khoury, R., Wang, W., Byrd, T.A., Pehek, E.A., Thacker, C., Zhu, X., Smith, M.A., Wilson-Delfosse, A.L., and Chen, S.G. (2010). LRRK2-mediated neurodegeneration and dysfunction of dopaminergic neurons in a *Caenorhabditis elegans* model of Parkinson’s disease. Neurobiology of Disease 40, 73–81. 10.1016/j.nbd.2010.04.002.

47. Cooper, J.F., Machiela, E., Dues, D.J., Spielbauer, K.K., Senchuk, M.M., and Van Raamsdonk, J.M. (2017). Activation of the mitochondrial unfolded protein response promotes longevity and dopamine neuron survival in Parkinson’s disease models. Sci Rep 7, 16441. 10.1038/s41598-017-16637-2.

48. Gitler, A.D., Chesi, A., Geddie, M.L., Strathearn, K.E., Hamamichi, S., Hill, K.J., Caldwell, K.A., Caldwell, G.A., Cooper, A.A., Rochet, J.-C., et al. (2009). α-Synuclein is part of a diverse and highly conserved interaction network that includes PARK9 and manganese toxicity. Nat Genet 41, 308–315. 10.1038/ng.300.

49. Desaeger, J.A., Rivera, M., Leighty, R., and Portillo, H. (2011). Effect of methomyl and oxamyl soil applications on early control of nematodes and insects. Pest Manag Sci 67, 507–513. 10.1002/ps.2058.

50. Clark, A.S., Huayta, J., Morton, K.S., Meyer, J.N., and San-Miguel, A. (2024). Morphological hallmarks of dopaminergic neurodegeneration are associated with altered neuron function in Caenorhabditis elegans. Neurotoxicology 100, 100–106. 10.1016/j.neuro.2023.12.005.

51. Morton, K.S., Hartman, J.H., Heffernan, N., Ryde, I.T., Kenny-Ganzert, I.W., Meng, L., Sherwood, D.R., and Meyer, J.N. (2023). Chronic high-sugar diet in adulthood protects Caenorhabditis elegans from 6-OHDA-induced dopaminergic neurodegeneration. BMC Biol 21, 252. 10.1186/s12915-023-01733-9.

52. Bradford, B.R., Whidden, E., Gervasio, E.D., Checchi, P.M., and Raley-Susman, K.M. (2020). Neonicotinoid-containing insecticide disruption of growth, locomotion, and fertility in Caenorhabditis elegans. PLoS One 15, e0238637. 10.1371/journal.pone.0238637.

53. Beilina, A., and Cookson, M.R. (2016). Genes associated with Parkinson’s disease: regulation of autophagy and beyond. J Neurochem 139 Suppl 1, 91–107. 10.1111/jnc.13266.

54. Risiglione, P., Leggio, L., Cubisino, S.A.M., Reina, S., Paternò, G., Marchetti, B., Magrì, A., Iraci, N., and Messina, A. (2020). High-Resolution Respirometry Reveals MPP+ Mitochondrial Toxicity Mechanism in a Cellular Model of Parkinson’s Disease. Int J Mol Sci 21, 7809. 10.3390/ijms21217809.

55. Reeve, A.K., Ludtmann, M.H., Angelova, P.R., Simcox, E.M., Horrocks, M.H., Klenerman, D., Gandhi, S., Turnbull, D.M., and Abramov, A.Y. (2015). Aggregated α-synuclein and complex I deficiency: exploration of their relationship in differentiated neurons. Cell Death Dis 6, e1820–e1820. 10.1038/cddis.2015.166.

56. Baker, M.J., Tatsuta, T., and Langer, T. (2011). Quality Control of Mitochondrial Proteostasis. Cold Spring Harb Perspect Biol 3, a007559. 10.1101/cshperspect.a007559.

57. Gao, A.W., El Alam, G., Lalou, A., Li, T.Y., Molenaars, M., Zhu, Y., Overmyer, K.A., Shishkova, E., Hof, K., Bou Sleiman, M., et al. (2022). Multi-omics analysis identifies essential regulators of mitochondrial stress response in two wild-type *C. elegans* strains. iScience 25, 103734. 10.1016/j.isci.2022.103734.

58. Greenwood, S.M., Mizielinska, S.M., Frenguelli, B.G., Harvey, J., and Connolly, C.N. (2007). Mitochondrial dysfunction and dendritic beading during neuronal toxicity. J Biol Chem 282, 26235–26244. 10.1074/jbc.M704488200.

59. Rego, A.C., and Oliveira, C.R. (2003). Mitochondrial dysfunction and reactive oxygen species in excitotoxicity and apoptosis: implications for the pathogenesis of neurodegenerative diseases. Neurochem Res 28, 1563–1574. 10.1023/a:1025682611389.

60. Sule, R.O., Condon, L., and Gomes, A.V. (2022). A Common Feature of Pesticides: Oxidative Stress-The Role of Oxidative Stress in Pesticide-Induced Toxicity. Oxid Med Cell Longev 2022, 5563759. 10.1155/2022/5563759.

61. Kondo, M., Yanase, S., Ishii, T., Hartman, P.S., Matsumoto, K., and Ishii, N. (2005). The p38 signal transduction pathway participates in the oxidative stress-mediated translocation of DAF-16 to *Caenorhabditis elegans* nuclei. Mechanisms of Ageing and Development 126, 642–647. 10.1016/j.mad.2004.11.012.

62. Aghayeva, U., Bhattacharya, A., and Hobert, O. (2020). A panel of fluorophore-tagged daf-16 alleles. microPublication Biology. 10.17912/micropub.biology.000210.

63. Ray, A., Martinez, B.A., Berkowitz, L.A., Caldwell, G.A., and Caldwell, K.A. (2014). Mitochondrial dysfunction, oxidative stress, and neurodegeneration elicited by a bacterial metabolite in a C. elegans Parkinson’s model. Cell Death Dis 5, e984–e984. 10.1038/cddis.2013.513.

64. Monti, D.A., Zabrecky, G., Kremens, D., Liang, T.-W., Wintering, N.A., Cai, J., Wei, X., Bazzan, A.J., Zhong, L., Bowen, B., et al. (2016). N-Acetyl Cysteine May Support Dopamine Neurons in Parkinson’s Disease: Preliminary Clinical and Cell Line Data. PLoS One 11, e0157602. 10.1371/journal.pone.0157602.

65. Xu, A., Shi, G., Liu, F., and Ge, B. (2013). Caenorhabditis elegans mom-4 is required for the activation of the p38 MAPK signaling pathway in the response to Pseudomonas aeruginosa infection. Protein Cell 4, 53–61. 10.1007/s13238-012-2080-z.

66. Iba, M., Kim, C., Kwon, S., Szabo, M., Horan-Portelance, L., Peer, C.J., Figg, W.D., Reed, X., Ding, J., Lee, S.-J., et al. (2023). Inhibition of p38α MAPK restores neuronal p38γ MAPK and ameliorates synaptic degeneration in a mouse model of DLB/PD. Sci Transl Med 15, eabq6089. 10.1126/scitranslmed.abq6089.

67. Chen, J., Li, M., Zhou, X., Xie, A., Cai, Z., Fu, C., Peng, Y., Zhang, H., and Liu, L. (2021). Rotenone-Induced Neurodegeneration Is Enabled by a p38-Parkin-ROS Signaling Feedback Loop. J. Agric. Food Chem. 69, 13942–13952. 10.1021/acs.jafc.1c04190.

68. Chikka, M.R., Anbalagan, C., Dvorak, K., Dombeck, K., and Prahlad, V. (2016). The Mitochondria-Regulated Immune Pathway Activated in the C. elegans Intestine Is Neuroprotective. Cell Rep 16, 2399–2414. 10.1016/j.celrep.2016.07.077.

69. Vérièpe, J., Fossouo, L., and Parker, J.A. (2015). Neurodegeneration in *C. elegans* models of ALS requires TIR-1/Sarm1 immune pathway activation in neurons. Nature Communications 6, 7319. 10.1038/ncomms8319.

70. Prins, N.D., de Haan, W., Gardner, A., Blackburn, K., Chu, H.-M., Galvin, J.E., and Alam, J.J. (2024). Phase 2A Learnings Incorporated into RewinD-LB, a Phase 2B Clinical Trial of Neflamapimod in Dementia with Lewy Bodies. J Prev Alzheimers Dis 11, 549–557. 10.14283/jpad.2024.36.

71. Yuan, W., Weaver, Y.M., Earnest, S., Taylor, C.A., Cobb, M.H., and Weaver, B.P. (2023). Modulating p38 MAPK signaling by proteostasis mechanisms supports tissue integrity during growth and aging. Nat Commun 14, 4543. 10.1038/s41467-023-40317-7.

72. Duarte-Jurado, A.P., Gopar-Cuevas, Y., Saucedo-Cardenas, O., Loera-Arias, M. de J., Montes-de-Oca-Luna, R., Garcia-Garcia, A., and Rodriguez-Rocha, H. (2021). Antioxidant Therapeutics in Parkinson’s Disease: Current Challenges and Opportunities. Antioxidants (Basel) 10, 453. 10.3390/antiox10030453.

73. Zuurbier, K.R., Solano Fonseca, R., Arneaud, S.L.B., Tatge, L., Otuzoglu, G., Wall, J.M., and Douglas, P.M. (2024). Cytosolic dopamine determines hypersensitivity to blunt force trauma. iScience 27, 110094. 10.1016/j.isci.2024.110094.

